# Systematic Analyses of Autosomal Recombination Rates from the 1000 Genomes Project Uncovers the Global Recombination Landscape in Humans

**DOI:** 10.1101/246702

**Authors:** Shivakumara Manu, Kshitish K Acharya, Saravanamuthu Thiyagarajan

## Abstract

**Background:** Meiotic recombination plays an important role in evolution by shuffling different alleles along the chromosomes, thus generating the genetic diversity across generations that is vital for adaptation. The plasticity of recombination rates and presence of hotspots of recombination along the genome has attracted much attention over two decades due to their contribution to the evolution of the genome. Yet, the variation in genome-wide recombination landscape and the differences in the location and strength of hotspots across worldwide human populations remains little explored.

**Results:** We make use of the untapped linkage disequilibrium (LD) based genetic maps from the 1000 Genomes Project (1KGP) to perform in-depth analyses of finescale variation in the autosomal recombination rates across 20 human populations to uncover the global recombination landscape. We have generated a detailed map of human recombination landscape comprising of a comprehensive set of 88,841 putative hotspots and 80,129 coldspots with their respective strengths across populations, about 2/3rd of which were previously unknown. We have validated and assessed the number of historical putative hotspots derived from the patterns of LD that are currently active in the contemporary populations using a recently published high-resolution pedigree-based genetic map, constructed and refined using 3.38 million crossovers from various populations. For the first time, we provide statistics regarding the conserved, shared, and unique hotspots across all the populations studied.

**Conclusions:** Our analysis yields clusters of continental groups, reflecting their shared ancestry and genetic similarities in the recombination rates that are linked to the migratory and evolutionary histories of the populations. We provide the genomic locations and strengths of hotspots and coldspots across all the populations studied which are a valuable set of resources arising out our analyses of 1KGP data. The findings are of great importance for further research on human hotspots as we approach the dusk of retiring HapMap-based resources.

## BACKGROUND

Genetic variability is primarily driven by mutations that create ‘variants of each gene’ (alleles). But meiotic recombinations provide an important means for shuffling the alleles of different genes along the chromosomes and thus enhance the genetic diversity across generations in the population. Recombination of various alleles across the genome permit removal of combinations that may negatively affect the carrying individual on one hand and provide a chance for generating beneficial combinations on the other. This additional mechanism of enhancing the overall genetic diversity within populations is favoured by natural selection as it enables sexually reproducing organisms to evolve more effectively and adapt to the changing environmental conditions [1]. It is also an obligatory feature of meiosis in most of the eukaryotes, since at least one crossover is essential for proper segregation of homologous pair of chromosomes [2]. Studying the sites of frequent recombinations, often referred to as ‘hotspots’, is important for understanding the evolutionary trends of a sexually reproducing species. Analysis of genomic data from ancient and contemporary human populations have already provided tremendous insights into the origin, migration, and expansion of our species (reviewed in [3]). According to a recent report [4], a subset of human populations first ventured out of Africa into other continents about 120,000 years ago. They must have encountered various selective pressures such as different climatic conditions, change in the diet, exposure to new pathogens, etc., and adapted differently to the local environments where the populations settled over thousands of years after migration. Under the varying influence of evolutionary forces, the landscape of recombination could have evolved differently across the populations. Establishing the similarities and differences in the location as well as the frequency of utilization of recombination hotspots across worldwide populations would be particularly interesting in terms of uncovering the global recombination landscape in humans.

Cytogenetic and linkage studies in the model organisms provided clues regarding the existence of hotspots of recombination at low-resolution [5]. Several efforts have been made to explore the location of hotspots as well as their frequency of utilization. Sperm-typing studies in humans characterised several of them at a high resolution indicating the presence of 1-2 kilobase (kb) long hotspots with strengths varying from ten to several hundred folds in comparison to the genome average [6]. However, this could not be applied to characterize the hotspots on a genome-wide scale due to the limitations of the sperm-typing method. Subsequent statistical analysis of population genetic data affirmed the substantial fine-scale variation in the recombination rates across the human genome [7]. Further analysis of linkage disequilibrium (LD) based fine-scale genetic maps developed using the polymorphism data of four representative populations from dense Perlegen SNP arrays led to the discovery of about 25,000 hotspots and their sequence contexts [8]. Later, the list was updated to about 33,000 hotspots and a degenerate 13-mer motif (CCNCCNTNNCCNC) that was enriched in 40% of the hotspots was reported using additional polymorphism data from the HapMap project phase 2 [9]. These developments in the area were followed up with the comparisons of the human hotspot locations with those in chimpanzees. The study [10], surprisingly, revealed no conservation. This difference was eventually accounted to a highly polymorphic protein called PRDM9 which was found to bind to the previously discovered 13-mer motif through its array of zinc-fingers [11, 12, 13]. Soon, a set of hotspots active only in the Africans [14] and another set of hotspots unique to each gender were unveiled [15]. These rapid advances spurred a wave of theoretical models and simulations, which tried to connect the empirical evidence and explain the evolutionary nature of hotspots [16, 17, 18]. Clearly, the recombination hotspots are fluidic elements of the genome. The plasticity and evolution of recombination rates along the genome are currently being widely researched, as the accumulating empirical evidence has unveiled the possibilities of a complex regulation of meiotic recombination operating at multiple levels (reviewed in [19]).

The initial broad-scale comparisons of patterns of LD and estimated recombination rates across the populations of Han Chinese, African Americans, and Americans of European descent showed no significant difference [20]. But, genome-wide and/or locus-specific influences of selection pressures may vary depending on the ecosystems of populations. Hence, it would not be surprising if the recombination rates along the human genome have been moulded differently across geographical locations [21]. In fact, a higher resolution study in a restricted 1 Mb segment of chromosome 22q across worldwide populations indicated the presence of extensive heterogeneity in the fine-scale recombination rate estimates [22]. Another pair of studies took advantage of the recent admixture in the African Americans and African Caribbeans to discover a set of hotspots active in the people of West African ancestry that are nearly inactive in the Europeans [14, 23]. From these studies, it was clear that the data from representative populations alone is not sufficient to acknowledge the entire variation in the recombination landscape across geographical regions because substantial variation exists even among the populations within a geographical region. The actual extent of such differences on a genome-wide scale remains to be established. A thorough analysis of fine-scale variation in the genome-wide recombination rates across multiple populations within and across continents is thus required to fully appreciate and understand the plasticity of the recombination landscape in humans. It can also help to begin research in new directions such as the potential role of allele-combinations in dictating certain phenotypes, disease susceptibilities, and drug-responses in populations. A comprehensive fine-scale analysis was delimited by the unavailability of genome-wide recombination rates for a wide range of populations, and computationally expensive methods for detecting and validating hotspots which were not feasible to apply on a genome-wide scale on multiple populations [24].

Advances in genomic technologies have led to the population-scale sequencing projects such as the 1000 Genomes Project [25] which has cataloged the human genetic variation at unprecedented depths. It also provided LD-based autosomal recombination rates for 20 worldwide populations estimated using coalescent methods through the LDhat program [26]. We capitalize on the availability of such extensive data for the first time to analyse the global recombination landscape in humans. Through alternative methods, we derive putative hotspots and coldspots from each population, validate them, and perform a comparative analysis across populations. We provide a comprehensive set of putative recombination hotspots and coldspots with their respective strengths across populations (Additional Files 1 and 2). Since the patterns of LD are influenced by recombination events accumulated over thousands of years through many generations, they reveal the historical hotspots along the genome [27]. Due to the rapid turnover of recombination hotspots, all of them might not be active in the contemporary populations. Also, some of the hotspot-like peaks might just be the noise from the coalescent estimator. Hence, we have also validated and evaluated the fraction of historical putative hotspots currently active in the extant populations utilizing a recently reported pedigree-based refined genetic map constructed using 3.38 million crossovers captured over 100,000 meiotic events from various populations [28]. For the first time, we highlight the conserved, shared, and unique validated hotspots present within and across geographical regions. Our analysis is useful in the backdrop of a previous research which established that by integrating various population-specific LD maps, the variation in individual hotspots can be captured and a global map of hotspots that can predict almost all the present-day crossovers can be developed [29]. Our hotspot map (Additional File 1) can also be used to ascertain gene conversion and de novo mutations arising during recombination [30]. Our study provides valuable resources for further research on characterizing these genomic regions to answer the long-standing questions about the dynamics and genetic basis of recombination at hotspots.

## RESULTS

### Global recombination landscape reflects migratory and evolutionary histories of the populations

We first established the fine-scale variation in the LD-based high-resolution genetic maps between the populations through holistic comparisons of autosomal recombination rates at various chromosomal positions. We achieved this by first rendering the 1KGP maps directly comparable through interpolation of missing SNP markers and then deriving the pairwise Pearson correlation coefficient. The extent of similarity expressed as Pearson correlation coefficients ranged from 0.63 to 0.89 indicating a linear relationship between the population-specific recombination landscapes. The pairwise correlation matrix of all the 20 populations taken up in this study (Supplementary Table 1, Additional File 3) establishes the amount of variation in the recombination landscapes among them (Supplementary Table 3, Additional File 3). To infer a global pattern of the variation in the recombination landscape, we clustered the pairwise correlation data. It can be seen that the populations that belong to a particular continent tend to cluster together (Figure 1). The African clade stands out from the other clades indicating minimum similarity of the African populations with every other geographical population considered. This observation is in line with the out-of-Africa migration theory and further expansion of human populations (reviewed in [3]). The recombination rates show a higher similarity between the populations within a geographical region than the distant populations. For example, the European populations show the highest homogeneity within a continent with a narrow range of correlation coefficients between 0.84 to 0.89, while the African populations have a range of comparatively lower correlation coefficients between 0.63 to 0.78 when compared across continents. The recombination landscapes of the Latin American populations are more similar to the European populations, which is an expected pattern given a relatively recent European ancestry. A seemingly contradicting observation is an Indian population clustering along with the European populations rather than the East Asian populations considering their geographical proximity. It should, however, be noted that the Indian populations are poorly represented in the current study and that a significant portion of Indians, particularly those living in the Northern parts including the Gujarat state, may have an ancient origin from Europeans, Central Asians, and Middle Easterners [31]. Further, the clades of East Asians, Americas, and the Europeans can be seen branching out from the common African ancestry (Figure 1). Thus, it is obvious that the clusters of continental groups and subpopulation structures reflect their shared ancestry and the variation in the global recombination landscape follows a pattern is linked to the migratory and evolutionary histories of the populations.

**Figure 1:**
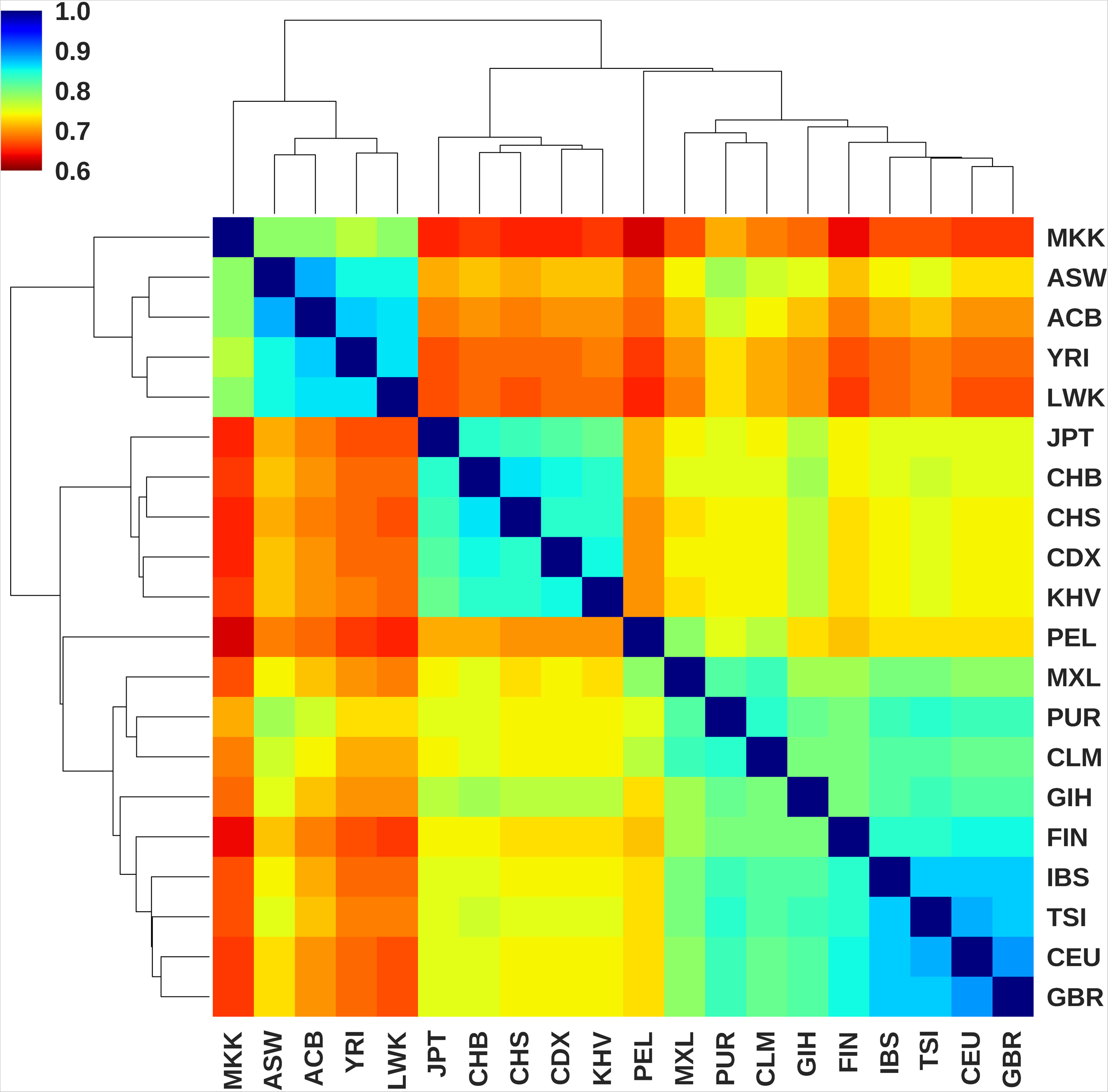
Cluster map displaying the correlation pattern of global recombination landscape from 20 human populations. Labels on the axes represent the 1KGP population codes (Supplementary Table 1, Additional File 3). The pairwise Pearson correlation coefficient matrix was clustered by hierarchical agglomerative method using the Euclidean distance metric and Unweighted Pair Group Method with Arithmetic Mean (UPGMA) algorithm. The clusters represent populations with similar autosomal recombination landscapes. The colour space of the heatmap represents the pairwise Pearson correlation coefficient between the fine-scale recombination rates between populations ranging from 0.6 to 1. See Supplementary Table 3, Additional File 3 for actual values and the results for data interpretations.

### Combined genetic map reveals putative hotspots and coldspots

We generated a cosmopolitan genetic map to capture the variation in the fine-scale recombination rates across populations at the level of individual hotspots and coldspots of recombination. We combined all the population-specific LD-based interpolated maps by averaging the genetic map units for all the SNP markers while deriving this map. We then identified putative hotspots and coldspots from the combined map using an adopted peak identification algorithm [29]. We found a total of 88,841 putative hotspots and 80,129 putative coldspots on the autosomes based on historical frequency of activity in one or more populations as they were inferred using LD-based recombination rates. We have provided the genomic locations of this comprehensive set of putative hotspots and coldspots along with their respective strengths across populations (Additional Files 1 and 2). We compared the genomic locations of these putative hotspots with the previously identified hotspots (N=32858) in phase 2 of the HapMap project [9]. We found that almost all of the LDhot-based HapMap phase 2 hotspots were present in our putative list, while the remaining 2/3^rd^ of them were novel.

### Cross validation using Pedigree-based refined genetic map

We compared the genomic locations of putative hotspots and coldspots revealed by the coalescent-based recombination rates with the recently published pedigree-based high-resolution refined genetic map [28] to validate the nature of putative regions and to avoid the noise from the coalescent-based estimator. The pedigree-based recombination rates were originally generated from over 3.38 million crossovers from various populations (Supplementary Table 2, Additional File 3) compiled from several studies [28]. A total of 54,066 hotspots and 51,691 coldspots were validated from the putative list of hotspots and coldspots using the pedigree-based recombination rates. This validated list also represents the fraction of hotspots and coldspots that are currently active in the contemporary populations as the pedigree-based rates were developed utilizing a large number of crossovers observed in quartet families from various populations. More than half of this validated list of hotspots are novel which were previously not detected in phase 2 of the HapMap project [9]. The observations indicate that the unvalidated hotspots possess lower strengths than the validated hotspots while the unvalidated coldspots possess comparatively higher strengths than the validated coldspots (Figure 2 and Supplementary Table 4, Additional File 3). Also, the LD-based and pedigree-based cosmopolitan strengths of validated hotspots are in good agreement with each other with a Pearson correlation coefficient of 0.79. The distance between the validated cosmopolitan hotspots extremely varies between few tens to couple of hundred kilobases with a mean width of about 33 kb. It should be noted that the mean strengths and the average distance between hotspots increase when population-specific hotspots and coldspots are considered because the cosmopolitan strengths are averaged across 20 populations and the list contains an exhaustive set of hotspots comprising of conserved, shared, and unique hotspots from all the populations. The mean width of validated hotspots is 4.88 kb (Std. dev. 3.77 kb) and that of coldspots varies largely with a mean width of about 28 kb.

**Figure 2:**
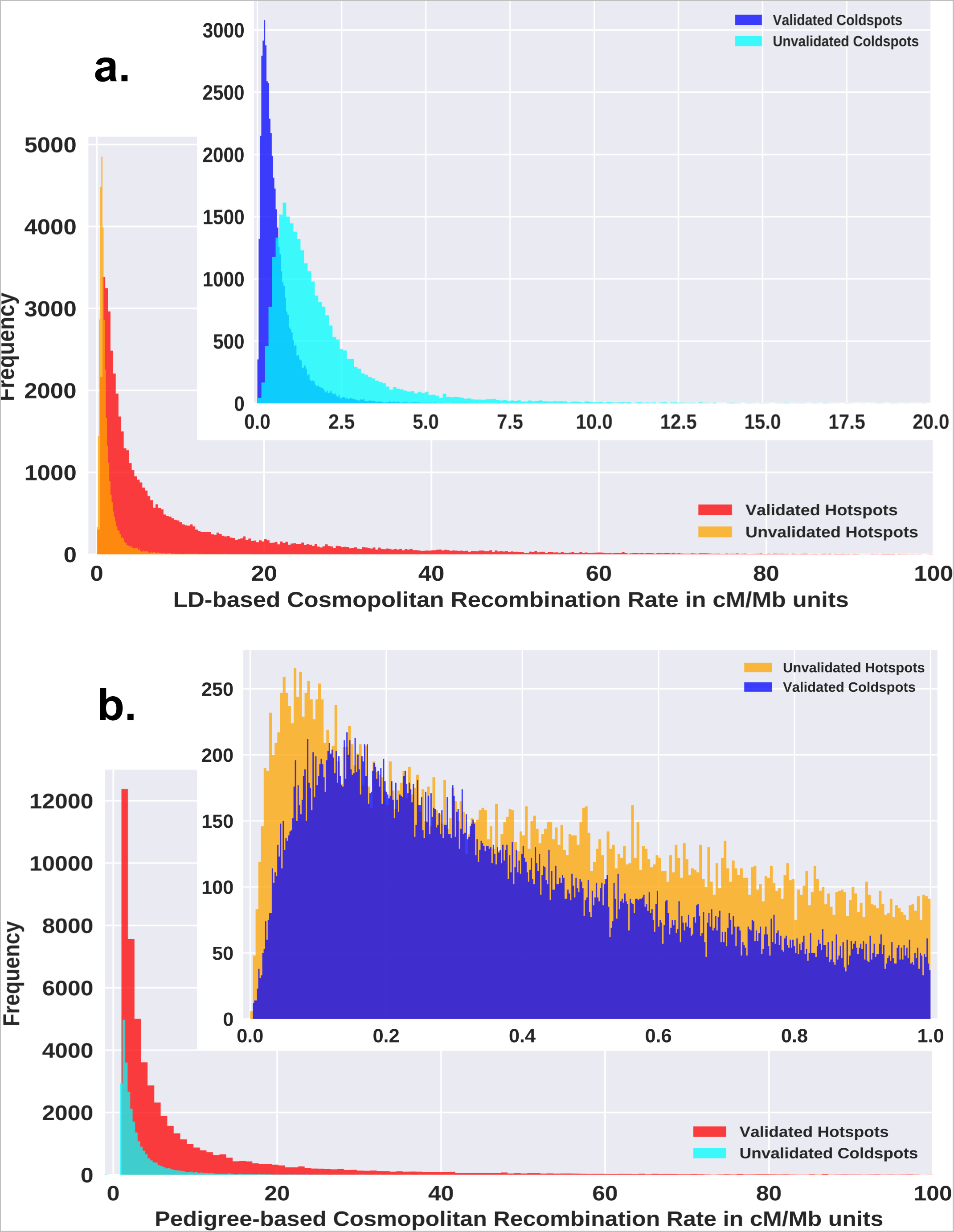
Relative frequencies of (a) LD-based and (b) Pedigree-based strengths of validated hotspots and coldspots averaged across all the studied populations (bin size=500). The validated hotspots possess higher strengths than the unvalidated hotspots and the validated coldspots possess lower strengths than unvalidated coldspots. See Supplementary Table 4, Additional File 3 for the mean and standard deviations of the distributions.

### Global distribution pattern of conserved, shared, and unique hotspots

We derived the number of shared hotspots between populations from the validated set of hotspots. The total number of hotspots in a population and the pairwise shared hotspots are tabulated in Supplementary Table 5, Additional File 3. A heat map of the shared hotspots between populations is depicted in Figure 3. African populations have the highest number of hotspots (about 46,500 to 48,250) reflecting the rapid decay in their LD that has been previously observed [32] while the East Asian populations share the least number of hotspots with other populations (about 37,000 to 40,000). The populations within a geographical region have similar shared hotspot profiles. We found 27,250 hotspots and 25,782 coldspots from the validated set to be conserved across all the 20 populations. Additionally, we detected a small number of validated hotspots that were unique to a super-population and active in one or more subpopulations. We found 1050 African, 302 American, 301 East Asian, 224 European, and 36 South Asian hotspots that were unique to the respective geographical regions, albeit many of them were weak. The distribution pattern of hotspots does not perfectly mirror the global correlation pattern of recombination landscape implying that the differences in the strengths of hotspots contribute substantially to the variation in the recombination landscape across populations along with the differences in the hotspot locations.

**Figure 3:**
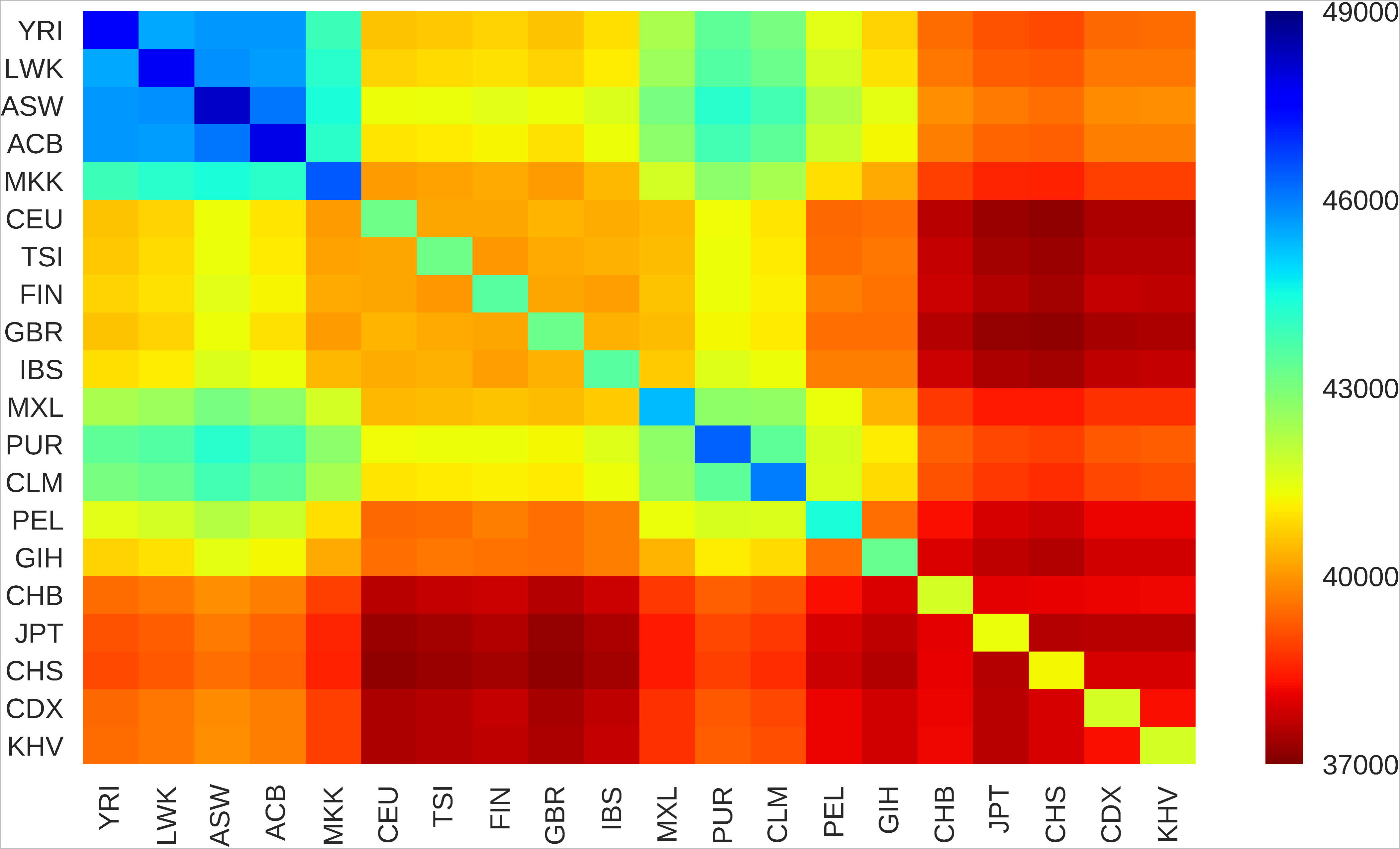
Heatmap of shared hotspots between populations computed from the validated set. Labels on the axes represent the 1KGP population codes(Supplementary Table 1, Additional File 3). The diagonal elements represent the total number of validated hotspots in the respective populations. The colour space of the heat map represents the number of shared hotspots between populations ranging from 37,000 to 49,000. See Supplementary Table 5, Additional File 3 for actual values and the results for data interpretations.

## DISCUSSION

The current understanding of the evolution of hotspots is based on the widely accepted red queen model [18]. In line with this model, a recent study [33] argued that the extant hotspots are active since the split of modern humans and Denisovans and are constantly evolving. The location and strength of recombination hotspots are under the influence of many evolutionary forces and the intricate interactions between them carve the landscape of recombination. It is potentially expected that the strength of these acting evolutionary forces varies between populations in a locus-specific manner or on a genome-wide basis and the recombination rates along the genome would be moulded accordingly [21]. The extent to which the recombination landscapes vary across populations was little known. The genome-wide dense genotype data and the fine-scale recombination rates between them furnished by the 1000 Genomes Project [25] has made it possible to compare the genome-wide recombination landscape across worldwide populations for the first time. We use this opportunity to capture the global recombination landscape in humans by thorough analyses of autosomal recombination rates and quantify the extent of similarities between populations. A minimum of 11% dissimilarity was observed even among the populations which are geographically nearby. A maximum of 37% dissimilarity was seen among distant populations belonging to different continents. We show that the clustering pattern of these similarity values reflects shared ancestry and it is linked to the migratory and evolutionary histories of the populations. We also identify 88,841 putative hotspots and 80,129 putative coldspots including most of the previously discovered hotspots (about 32,000) in the HapMap phase 2 project [9]. These hotspots were historically active in one or more populations since we obtained them by averaging all the population-specific genetic maps that were originally estimated by assessing the LD patterns through coalescent methods [26]. LD-based population-specific genetic maps are a superimposition of several influential allele-specific genetic maps and the contribution of each of them is proportional to the frequencies of the influential alleles in the population [34]. It implies that the hotspots derived from the sex-averaged and population-averaged genetic maps are the exhaustive set of hotspots and a fraction of them are used in any given individual. Basically, the choice of hotspots and their strength in an individual is a function of the underlying genetic makeup which is yet to be fully understood. Any genotype-phenotype mapping effort in this direction would require the usage data of the complete set of candidate hotspots from individuals with different genetic backgrounds which can be deduced through single-cell sequencing of sperms of individuals. It will be possible to make such large-scale efforts with the reduction in the cost and advancements in the throughput of single-cell sequencing technologies.

A key limitation of the LD-based approach used in our study is that the recombination rates are inferred using a population genetic model that does not take into account the effect of migration, natural selection, and varying effective population sizes on the patterns of LD [27]. There has been considerable debate over the effects of these phenomena that the elevated recombination rates at the hotspots might be exaggerated [35]. Although these maps can suggest the locations of potential hotspots, a separate method to test their significance is recommended to avoid the noise from the estimator. Currently used methods like LDhot [24] are computationally expensive, sensitive to the background window size and possess a moderate power of detection [36]. Here, we adopt a peak calling algorithm from a previously reported study [29] to identify putative regions coupled with the comparisons of their locations to a recently published high-resolution pedigree-based genetic map [28] constructed and refined from millions of crossovers across various populations to validate 54,066 hotspots and 51,691 coldspots. The pedigree-based genetic map had approached comparable levels of resolution to the LD-based maps through refinement using a Bayesian Markov Chain Monte Carlo procedure [28]. We also show that the LD-based and pedigree-based cosmopolitan strengths of the validated hotspots are in good agreement with each other with a Pearson correlation coefficient of 0.79. This amount of correlation between the recombination rates inferred from two different statistical methods is impressive, given the different proportions and compositions of the populations used in the original studies [25, 28] (Supplementary Tables 1 and 2, Additional File 3).

Further, for the first time, we also provide statistics about the conserved, shared, and unique hotspots across populations. The observed differences in the locations and strengths of the hotspots across populations might be due to the rapid turnover of hotspots under the influence of modifier loci and polymorphisms acting in cis or trans [18]. The highly evolving PRDM9 gene in the metazoans [37] is thought to be a major determinant of hotspots in many species [11, 12, 13]. Extensive variation is present in its tandem array of zinc-fingers and its motif binding properties within and across various species with over hundreds of alleles reported till date [38]. It will be interesting to see if the allele frequencies of different variants of PRDM9 among the studied populations correlate with the patterns of conserved, shared, and unique hotspots that we have furnished. While the polymorphism in this gene is one such known example, more are yet to be discovered because PRDM9 alone cannot account for all the variations observed and it is now known that several orders of birds and fishes as well as canids do not possess a fully functional copy of PRDM9 [39], yet successfully complete meiosis with recombinations.

The molecular mechanism of recognition of hotspots during meiosis has been an intriguing aspect for researchers. The 13-mer motif (CCNCCNTNNCCNC) is considered to be enriched in 40% of the hotspots [9] and recognised by the zinc-fingers of PRDM9 [11, 12, 13]. Although the details are not presented here, we did try to find any patterns associated with the hotspots. We used the discriminative search mode of the MEME program [40] with default parameters, and compared the top 100 validated hotspots with a control set of 100 validated coldspots of similar lengths. This top test set was selected based on the order of cosmopolitan LD-based strengths. We found multiple 10 to 30-mer motifs enriched in the selected hotspots including the 13-mer motif (CCNCCNTNNCCNC) and Poly A/T stretches. However, our search for these motifs using the FIMO [42] and MAST [43] programs, again using default parameters, showed that the number of the enriched patterns was similar in tested hotspots compared to the control set of coldspots. It should be noted that the 13-mer motif can be found to occur over 300,000 times in the human genome and is not exclusively present in the hotspots [35]. Although the red queen dynamics [18] between PRDM9 and its target motif seem to explain the empirical evidence concerning the hotspot turnover in evolutionary timescales, potential significance of the 13-mer motif remains questionable [35]. We noticed that these patterns were present inside the THE1, Alu, and LINE2 elements as reviewed elsewhere [41]. In general, there is a possibility that genome-wide repeat sequences could be important in defining hotspots. Hence, we also tried to find if any of the genome-wide repeats are exclusively associated with the same set of test sets using the GREAM tool [44] but did not find any statistically significant over-represented repeat elements within the sequences. The mechanism(s) of recognition of the hotspots in cells is perhaps more complex than expected and, currently, it may not be possible to study them using the commonly used algorithms that have been designed to explore signature patterns.

Finally, the development of faster and effective methods to infer recombination rates utilizing the whole genome variations and the usage of 1KGP phase 3 [25] polymorphisms in the latest genome build will increase the accuracy of estimates of hotspots, especially in low SNP density regions, low complexity regions, and gaps in the genome assembly.

## CONCLUSIONS

We have compared the autosomal recombination landscapes of 20 different worldwide populations and quantified the extent of similarities between them to reveal the global recombination landscape of human populations. Through clustering, we have shown that the differences in the recombination landscapes reflect shared ancestry and it is linked to the migratory and evolutionary histories of the populations. We have identified thousands of novel hotspots and coldspots that were active in one or more populations. We have also validated a large number of putative hotspots and coldspots using previously published pedigree-based recombination rates that were generated using millions of crossovers observed in various populations. We have also derived the conserved, shared, and unique hotspots in these populations and show that the African populations have the highest number of hotspots and the East Asian populations share the lowest number of hotspots with other populations. We have furnished the genomic locations as well as their strengths in each population for all the hotspots and coldspots. These valuable resources are useful in the further exploration of recombination hotspots and to deepen the understanding about their molecular basis as well as their evolutionarily transient nature.

## METHODS

### Collection of published genetic maps and hotspots

The linkage disequilibrium based recombination rates specific to 20 populations (Supplementary Table 1, Additional File 3) were downloaded from the official FTP site [45]. These rates were originally inferred by the coalescent-based LDhat program [26] utilizing the OMNI 2.5M phased genotype data from the 1000 Genomes Project samples [25]. The recently reported [28] high-resolution pedigree-based sex-averaged refined genetic map developed using millions of crossovers from various populations (Supplementary Table 2, Additional File 3) was obtained from the author’s GitHub repository [46]. The lifted-over HapMap phase 2 hotspots from a previously published work [47] were also included in this study. The genomic coordinates of all the utilised data in this study were in accordance with the GRCh37/hg19 build.

### Interpolation of 1KGP OMNI maps

The downloaded high-resolution genetic maps generated in the 1000 Genomes Project (1KGP) enumerate the population-specific recombination landscape along the autosomes at a very fine scale (Supplementary Figure 1, Additional File 4). They contain the physical coordinates of SNP markers in base pairs (bp) and their corresponding genetic distances in centimorgans (cM) with respect to the first SNP marker on the chromosome (Supplementary Table 6, Additional File 3). All the SNP markers that were used in the OMNI 2.5M genotyping array were not fully represented in all the population-specific maps. Hence, the physical coordinates of the missing SNP markers which were present in at least one of the maps were used to linearly interpolate the corresponding genetic distance in other maps (Supplementary Figure 2a, Additional File 4) using an in-house developed Python script. Extrapolation was not used for the SNP markers that were outside the range of a map as it would be error-prone. Instead, the genetic distance of a missing SNP marker which preceded the first SNP marker on the chromosome was set to zero and if it exceeded the range, it was set equal to the genetic distance of the last SNP marker on the chromosome. Interpolation ensured that there were no missing SNP markers and all the maps contained the same set of SNP markers with their respective genetic distances (Supplementary Figure 2b, Additional File 4).

### Pairwise correlation and clustering of the interpolated maps

The recombination rate between the adjacent SNP markers on the interpolated genetic maps was computed in terms of genetic distance per megabase of physical distance (cM/Mb) using a Python script. The arrays of recombination rates from all the chromosomes were concatenated together for each population retaining their order. The concatenated array of rates was compared across populations since they correspond to the same locations on the genome between the same set of SNP markers. Pearson correlation coefficient between the concatenated array of recombination rates was computed for all the pairwise combinations of populations to measure their linear relationships. Subsequently, hierarchical agglomerative clustering was performed on the pairwise correlation coefficients using the Euclidean distance metric and the Unweighted Pair Group Method with Arithmetic Mean (UPGMA) algorithm.

### Combining/averaging the interpolated 1KGP maps

To capture the overall trend in the recombination landscape, a cosmopolitan genetic map was generated by combining all the 20 population-specific interpolated 1KGP maps using a Python script. This was performed chromosome-wise by averaging the genetic map distance across all the populations for all the SNP markers. Any inconsistencies in the resulting averaged genetic map arising due to averaging were checked. After confirming the integrity of the averaged map, the recombination rate between the adjacent SNP markers of the cosmopolitan map was computed as described in the previous section (Supplementary Figure 3, Additional File 4).

### Deriving the putative hotspots and coldspots from the combined map

The combined cosmopolitan map generated by averaging all the population-specific interpolated 1KGP maps was utilised to derive the putative hotspots and coldspots. Hotspots appear as localised peaks in the linkage disequilibrium based recombination rate profile but a separate method is recommended to assess the significance of the peaks to avoid the noise from the estimator. The effectiveness of three different implementations of the commonly used LDhot [24] method to infer hotspots was recently assessed by Wall, J.D. et al. [36]. It cannot be currently applied on a genomewide scale across multiple populations due to its moderate power of detection, high sensitivity to the background window size, and computational complexity [36] (Supplementary Figure 4, Additional File 4). Hence, a peak identification algorithm was utilised to locate the candidate regions adopted from Khil, P.P. et al. [29]. This method relies on the accurate reconstruction of recombination rates by LDhat [26] and does not assess the significance of the localised peaks. The peak identification algorithm was modified to apply on the combined cosmopolitan genetic map as it was originally designed to analyse population-specific genetic maps. Hence, the initial peak detection step of the algorithm was modified while retaining the peak refinement step. The differences in the recombination rates between adjacent SNP markers were first calculated to locate the rising and falling points in the recombination rate profile of the combined cosmopolitan map. The block starting from the first rising point to the next rising point after a dip in the rate was computed which can potentially contain a peak. The block was then refined by fitting a Gaussian curve to the recombination rate profile. The refined start and end coordinates of the peak were then obtained by deriving the full-width half maximum (FWHM) of the Gaussian peak. The block was refined by cutting the profile at the mean rate of the block if the Gaussian fitting failed due to chaotic distributions. The process of block identification and refinement was then repeated by keeping the refined endpoint of the former peak as the start point of the latter block till the end of the last SNP marker on the chromosome. The consecutive refined regions without a gap in between were merged into one. Finally, the peaks present in the regions where the SNPs were sparsely typed (distance between typed SNPs > 50kb) and the peaks in the gaps of hg19 genome assembly were filtered. The approximate physical start and end positions of the candidate hotspots were then interpolated to obtain their respective genetic map coordinates in the combined cosmopolitan map as well as, in all the population-specific interpolated maps. The recombination rate (cM/Mb) between the start and end positions of the candidate hotspots were computed from all the maps to document their strengths across populations as described in the previous sections. The criteria to call a putative hotspot was that the computed strength of a candidate refined peak region should be above the average recombination rate of the human genome (1 cM/Mb) in one or more populations. The plateaus between putative hotspots were similarly evaluated for putative coldspots with the criteria that the computed strength should be below the genome average rate (1 cM/Mb) in one or more populations (Supplementary Figure 5a, Additional File 4).

### Validation of putative hotspots and coldspots using pedigree-based recombination rates

The putative hotspots and coldspots derived utilizing the linkage disequilibrium based recombination rates were validated by comparing their locations to the recently published pedigree-based population-averaged recombination rates [28]. This was possible mainly because this pedigree-based map was constructed by collecting over 3.38 million crossovers across different populations from the literature (Supplementary Table 2, Additional File 3) and had approached comparable levels of resolution to the LD-based maps through refinement using a Bayesian Markov Chain Monte Carlo procedure [28]. The start and end positions of the LD-based putative hotspots and coldspots were interpolated in the pedigree-based genetic map and their strength was computed with the same sex-averaged cut-off of 1 cM/Mb similarly as described in the previous section. This ensured that the hotspots and coldspots were verified using the recombination rates estimated from two different statistical methods for greater reliability (Supplementary Figure 5b, Additional File 4). A Pearson correlation coefficient between the LD-based and pedigree-based cosmopolitan strengths of validated hotspots was also computed to assess their agreement with each other.

### Assessing the hotspot usage across populations

The validated set of hotspots were used to derive the number of shared hotspots between all the pairwise combinations of populations. A hotspot was said to be shared between two populations if the strength of the hotspot exceeded the genome average recombination rate (1 cM/Mb) in both the populations. Additionally, a list of conserved hotspots which possessed strengths greater than the genome average recombination rate in all the 20 populations was derived. Similarly, a set of hotspots unique to a super population that were active only in one or more subpopulations were also detected (Supplementary Figure 6, Additional File 4).

## List of abbreviations

LD: Linkage Disequilibrium
1KGP: 1000 Genomes Project

## DECLARATIONS

### Ethics approval and consent to participate

Not Applicable

### Consent for publication

Not Applicable

## Availability of data and materials

All the in-house developed Python scripts, a pipeline, and all the raw-data required for reproducing the entire analyses are available at https://doi.org/10.5281/zenodo.1140907. The Hotspots and coldspots map resulting out of our analyses are also provided in hg19 coordinates.

## Author’s Contributions

SM conceived the project, collected the data, and wrote all the codes for analyses. SM, KA, and ST designed the study, analyses, and interpreted the results. SM wrote the manuscript with inputs from KA and ST. All the authors finalised and approved the manuscript.

## Acknowledgements

SM was partially supported through a Phi Psi Young Researcher Short Term Fellowship of the Institute of Bioinformatics and Applied Biotechnology (IBAB). We thank the DST-FIST support to IBAB vide project number SR/FST/LSI-536/2012. Institutional support to IBAB from the Department of IT, BT, S&T, GoK is greatly acknowledged.

## ADDITIONAL FILES

**Additional File 1.csv:** Human meiotic recombination hotspots map.

This file contains the genomic locations (hg19) and LD-based strengths of 88,841 hotspots across 20 populations. Additionally, information about the validation status, conservation across all the populations, averaged LD-based and pedigree-based cosmopolitan strengths are also provided for each hotspot.

**Additional File 2.csv:** Human meiotic recombination coldspots map.

This file contains the genomic locations (hg19) and LD-based strengths of 80,129 coldspots across 20 populations. Additionally, information about the validation status, conservation across all the populations, averaged LD-based and pedigree-based cosmopolitan strengths are also provided for each coldspot.

**Additional File 3.xlsx:** Supplementary Tables 1 to 6.

**Additional File 4.pdf:** Supplementary Figures 1 to 6.

